# Let-7b-5p differentially regulates human first trimester trophoblast migration and sFlt-1 through TLR7 and TLR8

**DOI:** 10.64898/2026.08.03.742516

**Authors:** Emily G Siegel, Lily C Salmeron, Vikki M Abrahams, Lubna Pal

**Affiliations:** Department of Obstetrics, Gynecology & Reproductive Sciences, Yale School of Medicine, New Haven, CT, USA

**Author notes:** Correspondence: Vikki M. Abrahams, PhD, Department of Obstetrics, Gynecology & Reproductive Sciences, Yale School of Medicine, 310 Cedar Street; LSOG 305C, New Haven, CT 06510, USA. **Additional Footnotes:** Authors contributed equally (Vikki M Abrahams, Lubna Pal).

**Keywords:** Angiogenic, MicroRNA, Migration, Preeclampsia, Toll-like Receptor, Trophoblast

## Abstract

**Introduction:** Preeclampsia is characterized by a pro-inflammatory, anti-migratory and anti-angiogenic placental phenotype. Impaired spiral artery remodeling stemming from trophoblast dysfunction is a key pathogenic mechanism. Little is known about the processes that govern trophoblast function normally and in preeclampsia. In preeclampsia, placental Let-7b-5p is reduced. The objectives of this study were to determine the normal function of Let-7b-5p in human trophoblast cells, to examine whether the ssRNA sensors, Toll-like receptor (TLR) 7 and/or TLR8 are mediators of trophoblast Let-7b-5p function, and whether disruption of this pathway promotes a preeclampsia-like phenotype in the trophoblast.

**Methods:** The human first trimester trophoblast cell line, Sw.71, was transfected with a Let-7b- 5p mimic, a Let-7b-5p inhibitor, or scramble control. Cells were treated with or without the TLR7 inhibitor IRS661 or the TLR8 inhibitor CUCPT9a. Trophoblast migration was measured using a two-chamber assay and interactions with human endometrial endothelial cells (HEECs) was measured using a 3D matrigel model. Trophoblast anti-angiogenic sFlt-1 release was measured by ELISA and sFLT1 mRNA measured by RT-qPCR.

**Results:** Transfection of trophoblast cells with a Let-7b-5p mimic elevated migration through activation of TLR7 and TLR8, while in a TLR7-dependent manner, the Let-7b-5p mimic negatively regulated sFlt-1 production. Furthermore, inhibition of trophoblast Let-7b-5p reduced migration, elevated FLT1 mRNA expression and sFlt-1 release, and reduced trophoblast-endometrial endothelial cell interactions.

**Conclusions:** This study highlights a role for TLR7/TLR8-activating Let-7b-5p in promoting normal trophoblast function and endothelial interactions and that disruption in this miR-driven signaling pathway may be relevant to processes driving a pre-eclamptic placental phenotype.

**Highlights:** Trophoblast migration is positively driven by Let-7b-5p activating TLR7 and TLR8

Let-7b-5p, via TLR7, negatively regulates trophoblast anti-angiogenic sFlt-1 production.

Inhibition of trophoblast Let-7b-5p reduces trophoblast migration and normal interactions with endometrial endothelial cells, while sFlt-1 production is elevated.

TLR7/TLR8-activating Let-7b-5p promotes normal trophoblast function and endothelial interactions and disruption in this miR-driven signaling pathway may promote a preeclamptic placental phenotype.

## 1. Introduction

Preeclampsia is a gestational hypertensive disorder that affects 2-8% of all pregnancies [1]. During normal placentation, cytotrophoblast cells differentiate into extravillous trophoblast cells as they invade deep into the maternal decidua, eventually invading the uterine spiral arteries and replacing the endothelial cells. This spiral artery remodeling by the invading trophoblast shifts the uterine and placental vasculature from a high resistance, high turbulence to a low-resistance, low pressure environment. However, in pregnancies that are destined for preeclampsia, the normal functionality of trophoblast is altered such that there is reduced cell migration and compromised arteriolar invasion by the invading extravillous trophoblast, as well as exaggerated release of inflammatory factors and anti-angiogenic factors, such as sFlt-1. These changes lead reduced remodeling of the spiral arterioles, rendering the placental bed a high-pressure, high-blood flow environment [2–5]. Despite, the pathophysiology of preeclampsia being established early in gestation, it is not clinically diagnosed until the second or third trimester, with delivery of the placenta being the only way to resolve symptoms. Due to its late diagnosis, it is important to investigate early indicators, such as biomarkers, that may determine a woman’s risk of developing preeclampsia.

One such potential biomarker is the microRNA (miR), Let-7b-5p. In human first trimester trophoblast cells, Let-7b-5p overexpression has been shown to drive cell invasion [6]. Although Let-7b-5p is upregulated in uncomplicated third trimester human placentas compared to first trimester samples [7], in patients with preeclampsia, placental and maternal circulating levels of Let-7b-5p are reduced [6, 8, 9]. A reduction in placental Let-7b-5p expression may, therefore, result in reduced trophoblast migration and invasion, which may negatively impact subsequent spiral artery remodeling; a hallmark of preeclampsia.

Let-7b-5p is one of a small family of GU-rich sequence-containing miRs that can act non-canonically by binding and activating the ssRNA sensors, Toll-like receptor (TLR) 7 and/or TLR8 [10]. Our group discovered another similar miR, miR-146a-3p, that drives human first trimester trophoblast inflammation through activation of TLR7 and TLR8 [11, 12]. We, therefore, questioned whether Let-7b-5p was able to regulate trophoblast migration, angiogenic factor production, and interactions with endometrial endothelial cells, also through TLR7 and/or TLR8 activation. Thus the objective of this study was to determine the normal function of Let-7b-5p in human first trimester trophoblast cells, whether TLR7 and TLR8 were involved, and whether disruption of this pathway would promote a preeclampsia-like trophoblast phenotype. Herein, we report that inhibition of Let-7b-5p and therein, the inhibition of TLR7 and TLR8, resulted in decreased trophoblast migration. Let-7b-5p inhibition also elevated trophoblast production of the anti-angiogenic factor sFlt-1 in a TLR7- and NFκB-dependent manner. Furthermore, using a three dimensional (3D) *in vitro* model of trophoblast invasion and subsequent interactions with endometrial endothelial cells similar to that seen in normal spiral arteriole transformation, inhibition of Let-7b-5p reduced trophoblast invasion and interactions with human endometrial endothelial cells. Finally we report that the reduced trophoblast-endothelial cross talk and/or endothelial angiogenesis may be mediated by trophoblast derived small extracellular vesicles (sEVs) with reduced Let-7b-5p expression. Together, this study highlights a role for TLR7/TLR8-activating Let-7b-5p in promoting normal trophoblast function and endothelial interactions and that disruption in this miR-driven signaling pathway may be relevant to processes driving a preeclamptic placental phenotype.

## 2. Materials and Methods

### 2.1. Cell lines

Studies were performed using the telomerase immortalized human first trimester extravillous trophoblast cell line, Sw.71 [13]. Trophoblast cells were cultured in full media comprising of Dulbecco’s modified Eagle’s medium/F-12 (DMEM/F-12; Life Technologies, Grand Island NY) with 10% fetal bovine serum (FBS) (HyClone, South Logan, UT, USA), 10mM MEM non-essential amino acids, 1mM sodium pyruvate, 10mM HEPES and 100nM penicillin/streptomycin (Life Technologies). For co-culture and angiogenic studies, the human endometrial endothelial (HEEC) cell line was used as previously described [14]. HEECs were cultured in EBM-2 growth media (Lonza, Basel, Switzerland) supplemented with 10% FBS. Cells were incubated at 37 C with 5% CO_2_.

### 2.2. Trophoblast cell transfections and treatments

Trophoblast cells were transfected using siPORT NeoFX (Invitrogen, Waltham, MA) with either a MirVana miR-Let-7b-5p mimic (MC11050; ThermoFisher Scientific, Waltham, MA) at 100nM, a miR mimic scramble control at 100nM (ThermoFisher Scientific), a MirVana miR-Let-7b-5p inhibitor (MH11050; ThermoFisher Scientific, Waltham, MA) at 50nM, or a miR inhibitor scramble control at 50nM (ThermoFisher Scientific). For some experiments, 1 hr post-transfection, cells were treated with or without the TLR7 inhibitor, IRS661 (5μM; made endotoxin-free by the Keck Core, Yale University) [15, 16], the TLR8 inhibitor, CUCPT9a (2.5μM) (Invivogen; San Diego, CA), or the NFκB inhibitor, BAY117085 (1μM) (Millipore-Sigma, Burlington, MA). In other experiments, trophoblast cells, in the absence of any transfection, were treated with either no treatment (NT), IRS661 at 5μM, CUCPT9a at 2.5 μM, the TLR7 agonist, R837 at 5μg/ml (Invivogen), or the TLR8 agonist, TL8-506 at 5μg/ml (Invivogen). All transfections and treatments were performed in serum-free OptiMEM (Life Technologies).

### 2.3. Trophoblast small extracellular vesicle (sEV) isolation

Following transfection, trophoblast cell culture supernatants were collected at 72 hrs and sEVs were isolated using ExoQuick-TC (Systems Biosciences LLC, Palo Alto, CA) per the manufacturer’s instructions. The isolated sEV pellet was either: 1) resuspended in 25μl of sterile phosphate buffered saline (PBS, Thermo Fisher Scientific, Waltham, MA) and 25uL of sterile nuclease free water (Invitrogen, Waltham, MA) for subsequent functional studies following nanoparticle tracking analysis (NTA) and quantification using a Zetaview (Particle Metrix, Ammersee, Germany) set to room temperature, a laser wavelength of 488nm, a sensitivity of 80, and a shutter speed of 100; or 2) directly lysed for RNA extraction using the SeraMir Exosome RNA Purification Column Kit (Systems Biosciences).

### 2.4. RNA isolation and RT-qPCR

Total cellular RNA was extracted using Trizol following manufacturer’s protocol. RNA concentrations were measured using a NanoDrop 2000 Microvolume Spectrophotometer (ThermoFisher Scientific; Waltham, MA). Cellular and sEV RNA expression of Let-7b-5p was measured using the Taqman miR Assay (Life Technologies; Carlsbad, CA) following the manufacturer’s protocol for reverse transcription and qPCR. Small nuclear RNA U6 was used as a reference gene for normalization. Trophoblast cellular mRNA expression of the FLT1 splice variant 2 (V2) was measured by RT-qPCR. This FLT1 splice variant (sFlt-1 i13) is one of the most commonly observed variant in the human placenta and gives rise to the secreted sFlt-1 protein [17]. FLT1 V2 mRNA was measured using the SuperScript II kit (Invitrogen; Waltham, MA) and the Kapa SYBR Fast qPCR kit (Sigma-Aldrich St. Louis, MO) and normalized to GAPDH. The primers used for detecting FLT1 V2 mRNA (NM_001159920.2) has a forward sequence of TTCCGAAGCAAGGTGTGACT, a reverse sequence of AGCCTTTTTGTTGCAGTGCTC, and an efficiency of 96.53%. The GAPDH primer has been previously reported [18]. RT-qPCR data was analyzed using the 2^-ΔΔCT^ method and reported as fold change (FC).

### 2.5. Protein isolation and Western blot analysis

Cell protein was extracted using lysis buffer containing phosphatase inhibitors (Cell Signaling) and concentrations measured using the Pierce BCA protein assay (ThermoFisher Scientific, Waltham, MA); 20μg of protein was resolved on a 10% SDS PAGE gel and then transferred onto a PVDF membrane (Bio-Rad). Membranes were probed for the following primary antibodies all from Cell Signaling Technology (Danvers, MA): phosphorylated (p)-p65 NFκB (#3033; 1:1000); total (t)-p65 NFκB (#8242; 1:1000); p-p38 MAPK (#9211;1:500); t-p38 MAPK (#9212; 1:1000); p-ERK (#9101; 1:1000); t-ERK (#4695; 1:1000); p-JNK (#9251; 1:1000); and t-JNK (#9252; 1:1000). Western blot images were captured using an Amersham Imager 680 (General Electric, Boston, MA). Image Studio Lite (Li-Cor Biosciences, Lincoln, NE) was used to perform semiquantitative densitometry. Fold changes were determined by normalizing phosphorylated protein levels against the paired total protein levels.

### 2.6. Trophoblast cell migration

A two-chamber colorimetric assay was used to measure trophoblast cell migration [19]. A 24 well tissue culture plate containing 800μl serum-free OptiMEM acted as the lower chamber of the assay. A cell culture well insert with an 8μm pore size membrane was used as the upper chamber (Millipore-Sigma, Burlington, MA, USA). Trophoblast cells (with or without transfection) suspended in 200μl serum-free OptiMEM, with or without treatments/inhibitors, were seeded into the upper chamber. After 24 or 48 hrs, spontaneous trophoblast migration across the membrane was measured using the QCM 24-well Colorimetric Cell Migration Assay and read in triplicate at 560nm using a Bio-Rad iMark Microplate Absorbance Reader (Hercules, CA, USA). Relative percent migration for each condition/treatment was calculated by comparing results to a 100% cell control.

### 2.7. Cell viability

Trophoblast and HEEC cell viability was determined using the CellTiter 96 viability assay (Promega, Madison, WI, USA), as previously described [19]. Cells (transfected or not) were seeded into wells of a 96-well plate in full media and cultured overnight. The media was then replaced with serum-free OptiMEM, and cells were incubated for an additional 3 hrs Cells were then treated or not and after 48 hrs, the CellTiter 96 substrate was added to all wells and incubated for 2 hrs at 37°C. Optical densities were read at 490 nm. All samples were assayed in triplicate, and cell viability was presented as a percentage of the untreated control.

### 2.8. Measurement of trophoblast secreted factors

Trophoblast supernatants were measured for IL-1β, IL-6, IL-8, IL-11, TNFα, VEGF, PlGF, sEndoglin, sFlt-1, and MMP14 by ELISA (R&D Systems, Minneapolis, MN) following manufacturer’s protocol. Cell protein lysates were also measured for MMP14 by ELISA.

### 2.9. Trophoblast-HEEC 3D co-culture assay

Studies to investigate trophoblast invasion and subsequent interactions with endometrial endothelial cells, similar to that seen in normal spiral artery transformation, were performed using a 3D *in vitro* system, as previously described [20]. HEECs were stained with the red-fluorescent linker dye, PKH-26 (Sigma, St. Louis, MO). HEECs were seeded into 24-well tissue culture plate (2x10^5^ cells/well) over undiluted reduced growth factor Matrigel (BD-Biosciences, San Jose, CA) and cultured in 250μl of supplemented EBM-2 growth media overnight until tube-like structures were observed. Media was removed and trophoblast cells, transfected with either a scramble control or a Let-7b-5p inhibitor, were stained with the green-fluorescent linker dye, PKH-67 (Sigma, St. Louis, MO) and then were seeded (1x10^5^ cells/well) in 500μl OptiMEM. The trophoblast-endothelial co-culture system was incubated for 24 hrs and then imaged by fluorescent microscopy (Keyence BZ-X800 Series) at 4x magnification using BZ-X Analyzer software. For each well, 5 images were collected and the % red and green fluorescence per field quantified using BZ-X Analyzer software and averaged.

### 2.10. HEEC 3D angiogenesis assay

An endothelial tube formation assay was used to measure HEEC angiogenesis [21, 22]. HEECs were seeded into 24-well tissue culture plates over undiluted reduced growth factor Matrigel together with trophoblast sEVs (1x10^9^/ml) isolated from trophoblast cells transfected with either a scramble control (Control sEV) or a Let-7b-5p inhibitor (Let-7b-5p inhib sEV). After 4hrs HEEC tubes were imaged by phase contrast microscopy (Keyence BZ-X800 Series) at 4x magnification using BZ-X Analyzer software. For each well, 5 images were collected and the number of HEEC tubes per image quantified and averaged.

### 2.11. Statistical analysis

Each experiment was performed at least three times. All analyses were performed in duplicate or triplicate. All data are reported as either mean ± standard error of the mean (SEM) or fold change ± standard error of the mean (SEM) of pooled experiments. The number of individual experiments from which data were pooled are indicated in figure legends as “n=”. Statistical significance was defined as *p*<0.05 and determined using Prism Software (GraphPad, La Jolla, CA, USA). Significance for normally distributed data was determined using either one-way analysis of variance (ANOVA) for multiple comparisons or a t-test. Significance for data of non-Gaussian distribution was determined using a non-parametric multiple comparison test or Wilcoxon matched pairs signed rank test.

## 3. Results

### 3.1. Let-7b-5p drives human first trimester trophoblast cell migration through activation of TLR7 and TLR8

Compared to trophoblast cells transfected with a scramble control, cells transfected with a Let-7b-5p mimic had significantly increased Let-7b-5p expression by 300.4 ± 55.3-fold after 24 hrs, and by 82.6 ± 29.2-fold after 72 hrs (Figure 1A). Conversely, transfection of trophoblast cells with a Let-7b-5p inhibitor significantly reduced basal Let-7b-5p expression by 85.0 ± 3.2% after 24 hrs, and by 48.4 ± 9.9% after 72 hrs, when compared to the scramble control (Figure 1A). Transfection of trophoblast cells with either the Let-7b-5p mimic or the Let-7b-5p inhibitor had no effect on cell viability compared to the scramble controls (Figure 1B). Overexpression of Let-7b-5p through mimic transfection significantly elevated trophoblast migration by 1.6 ± 0.1-fold when compared to the control, while Let-7b-5p inhibition significantly reduced trophoblast migration by 27.3 ± 7.2% (Figure 1C). To determine whether basally expressed Let-7b-5p was able to drive trophoblast migration through TLR7 and/or TLR8 activation, cells were treated with either a TLR7 or TLR8 receptor inhibitor or agonist. As shown in Figure 1D, treatment of trophoblast cells with the TLR7 inhibitor, IRS661, or the TLR8 inhibitor, CUCPT9a significantly reduced migration by 33.7 ± 6.9% and 29.5 ± 5.2%, respectively compared to the no treatment (NT) control. Conversely, compared to the NT control, only the TLR7 agonist, R837, significantly elevated trophoblast migration by 1.2 ± 0.1-fold (Figure 1E). The TLR8 agonist, TL8-506, had no effect on trophoblast migration (Figure 1E). Despite, a lack of effect of the TLR8 agonist on trophoblast migration, the increased trophoblast migration in response to the Let-7b-5p mimic was significantly reduced by the presence of either the TLR7 inhibitor, IRS661, or the TLR8 inhibitor, CUCPT9a demonstrating that Let-7b-5p was indeed driving trophoblast migration through both TLR7 and TLR8 activation (Figure 1F). Neither the TLR7 inhibitor nor the TLR8 inhibitor had any effect on trophoblast cell viability (Figure 1G).

**Figure 1.**
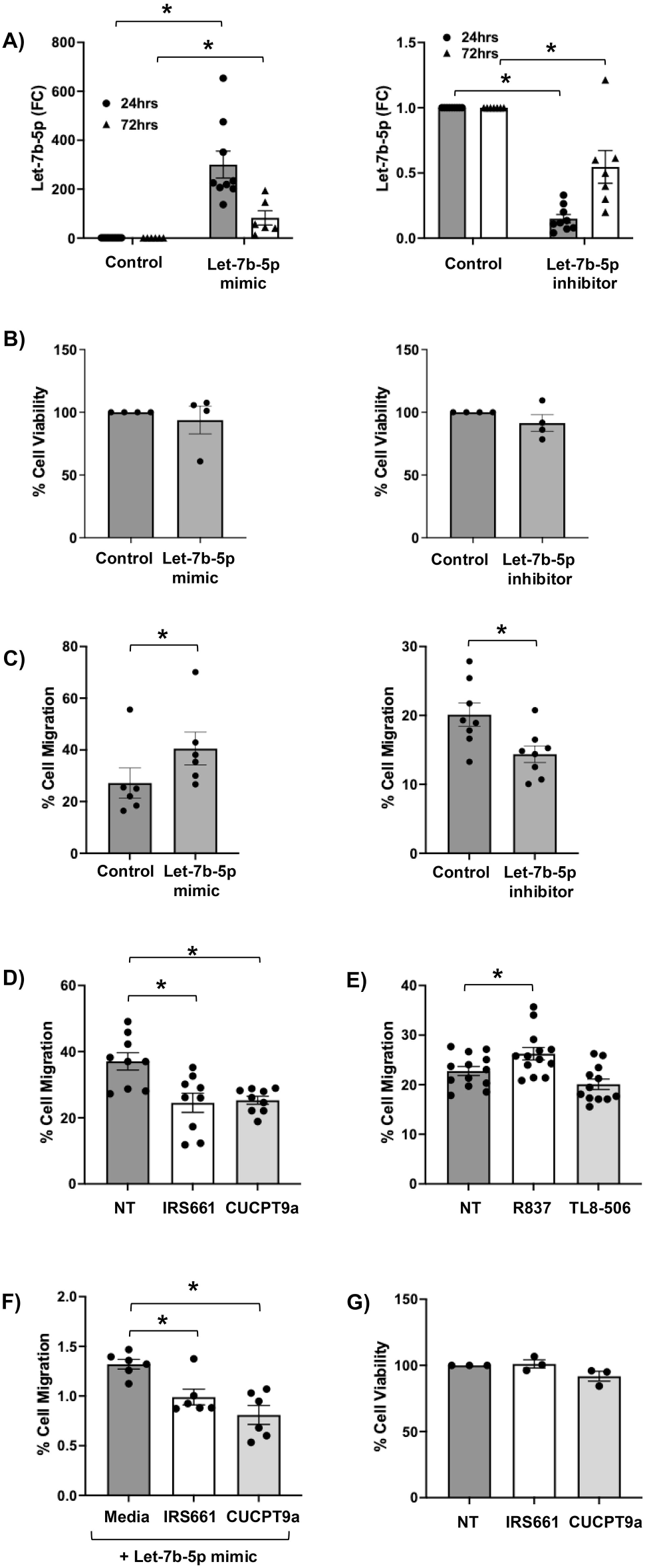
Let-7b-5p drives human first trimester trophoblast cell migration through activation of TLR7 and TLR8. (A-C) Trophoblast cells were transfected with a scramble control (control), a Let-7b-5p mimic or a Let-7b-5p inhibitor. (A) After 24 hrs and 72 hrs, Let-7b-5p expression was measured by RT-qPCR and expressed as fold change (FC) relative to the control (n=6-9; \**p*<0.05). (C) After 48 hrs trophoblast cell viability was measured (n=4). (C) After 48 hrs trophoblast cell migration was measured (n=6-8; \**p*<0.05). (D-E) Trophoblast cells were treated: (D) for 24 hrs with no treatment (NT), the TLR7 inhibitor, IRS661, or the TLR8 inhibitor, CUCPT9a (n=9; \**p*<0.05), or (E) for 48hrs with NT, the TLR7 agonist, R837, or the TLR8 agonist TL8-506 (n=14; \**p*<0.05), after which cell migration was measured. (F) Trophoblast cells were transfected with a scramble control or a Let-7b-5p mimic and then treated with either media, IRS661 or CUCPT9a. After 24 hrs, cell migration was measured and data are shown as the fold change (FC) of trophoblast migration under Let-7b-5p mimic conditions relative to the scramble control (n=6; \**p*<0.05). (G) Trophoblast cells were treated with NT, IRS661 or CUCPT9a and after 24 hrs, cell viability was measured (n=3).

### 3.2. Human first trimester trophoblast anti-angiogenic sFlt-1 production is negatively regulated by Let-7b-5p activating TLR7

To further explore how Let-7b-5p might be regulating trophoblast function, we tested its effect on the release of the anti-angiogenic factor, sFlt-1, which like trophoblast migration, is also altered in preeclampsia. Compared to trophoblast cells transfected with a scramble control, cells transfected with a Let-7b-5p mimic secreted 28.1 ± 5.9% less sFlt-1, and conversely, the presence of the Let-7b-5p inhibitor significantly elevated trophoblast sFlt-1 release by 1.3 ± 0.1- fold (Figure 2A). Since sFlt-1 is generated either through expression of an alternative splice variant or through proteolytic cleavage by the protease, MMP14 [23], we explored both levels of regulation in the trophoblast by Let-7b-5p. As shown in Figure 2B, the Let-7b-5p inhibitor significantly elevated trophoblast mRNA expression of the FLT1 splice variant 2, which gives rise to sFlt-1 (Figure 2B). In contrast, intracellular levels of trophoblast MMP14 protein levels were unchanged by Let-7b-5p inhibition (Figure 2C). Secreted MMP14 was mostly under the detection limit of the ELISA (data not shown). Similar to the Let-7b-5p mimic, treatment of trophoblast cells with the TLR7 agonist, R837, significantly decreased sFlt-1 release by 50.1 ± 3.8%, while the TLR8 agonist, TL8-506 had no effect (Figure 2D). As shown in Figure 2E, the reduced trophoblast sFlt-1 release in response to Let-7b-5p overexpression using the miR mimic was reversed back to baseline by the TLR7 inhibitor, IRS661, but not by the TLR8 inhibitor, CUCPT9a, demonstrating that the negative regulation of sFlt-1 by Let-7b-5p was mediated through the miR activating TLR7.

**Figure 2.**
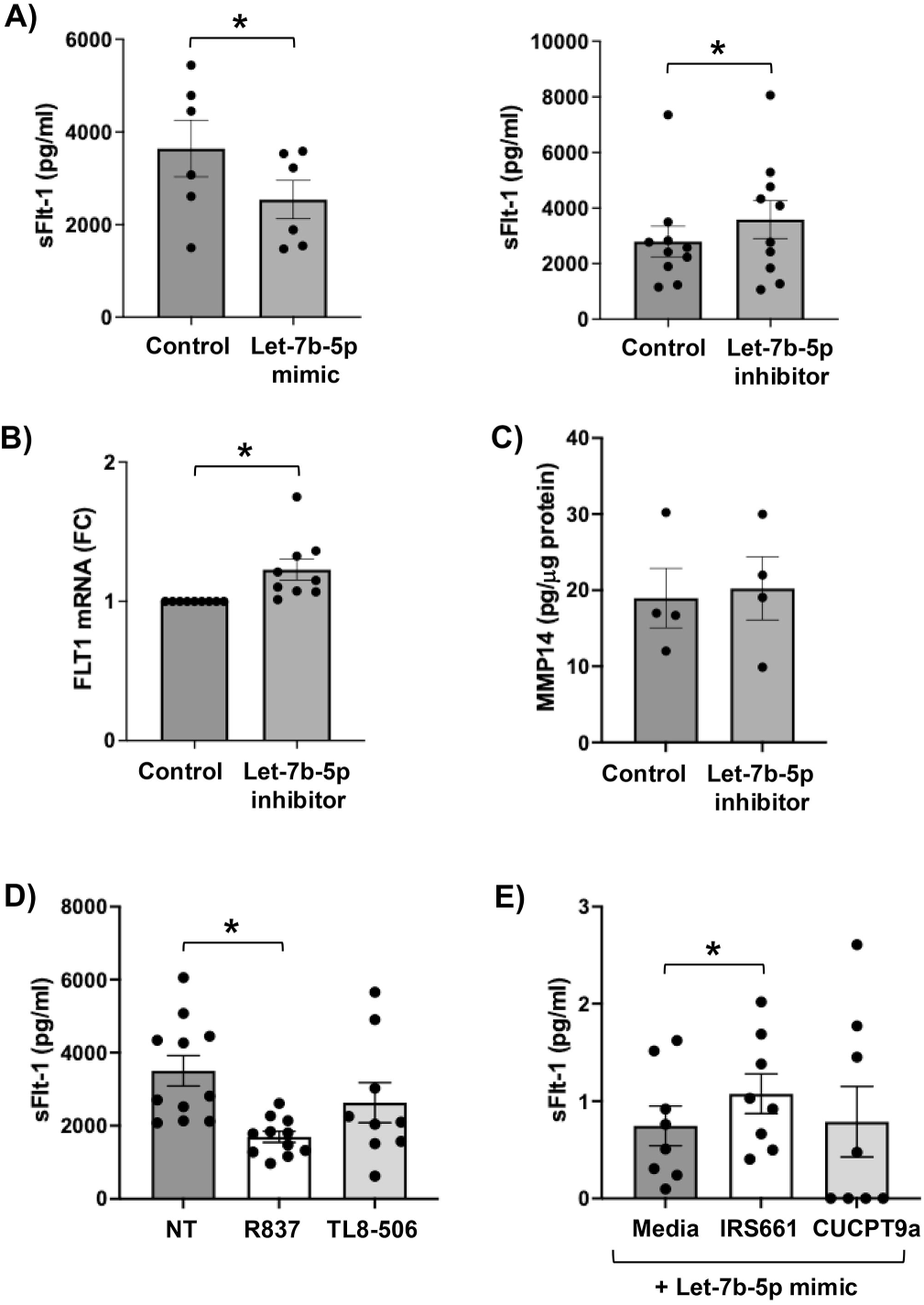
Human first trimester trophoblast anti-angiogenic sFlt-1 is negatively regulated by Let-7b-5p via TLR7 activation. (A-C) Trophoblast cells were transfected with a scramble control (control), a Let-7b-5p mimic (n=6) or a Let-7b-5p inhibitor (n=4-10). After 72 hrs, (A) supernatants were measured for sFlt-1 by ELISA (\**p*<0.05), (B) cellular RNA was measured for FLT1 mRNA levels by RT-qPCR using GAPDH as a housekeeping gene for normalization (\**p*<0.05), and (C) cell lysates were measure for MMP14 protein by ELISA. (D) Trophoblast cells were treated for 72 hrs with NT, R837, or TL8-506 (n=11), after which sFlt-1 secretion was measured by ELISA (\**p*<0.05). (E) Trophoblast cells were transfected with a scramble control or a Let-7b-5p mimic and then treated with either media, IRS661 or CUCPT9a. After 72 hrs, supernatants were measured for sFlt-1 by ELISA and data is shown as the fold change (FC) of sFlt-1 release relative to the scramble control (n=8;*\*p*<0.05).

### 3.3. Human first trimester trophoblast secreted factors that are not regulated by Let-7b-5p

Since we found that Let-7b-5p regulated sFlt-1 release by the trophoblast, we sought to investigate if other trophoblast secreted factors that might be dysregulated in preeclampsia were also altered. Thus, we tested the effects of the Let-7b-5p mimic and Let-7b-5p inhibitor on trophoblast secretion of the inflammatory factors IL-8, IL-1β and TNFα; the pro-migratory factors IL-6 and IL-11; and the angiogenic factors VEGF, sEndoglin and PlGF. As shown in Figure 3, neither the Let-7b-5p mimic nor the Let-7b-5p inhibitor had any effect on the ability of trophoblast cells to secrete (A) IL-8, (B) IL-1β, (C) TNFα, (D) IL-6, (E) IL-11, (F) VEGF, or (G) sEndoglin. PlGF levels were mostly undetectable (data not shown).

**Figure 3.**
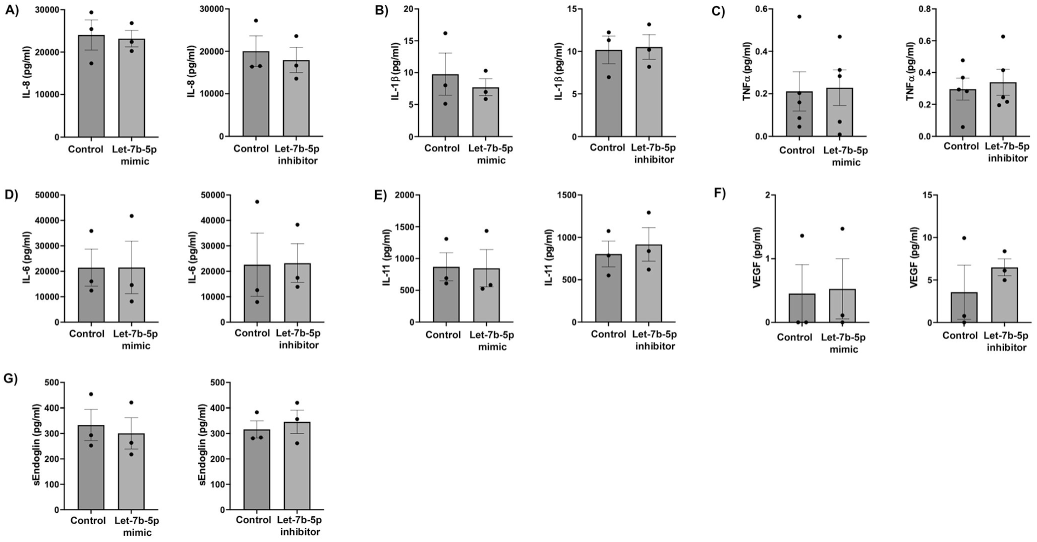
Human first trimester trophoblast secreted factors that are not regulated by Let- 7b-5p. Trophoblast cells were transfected with a scramble control (control), a Let-7b-5p mimic or a Let-7b-5p inhibitor (n=3). After 72 hrs cell supernatants were measured by ELISA for: (A) pro-inflammatory IL-8; (B) pro-inflammatory IL-1β; (C) pro-inflammatory TNFα; (D) pro-migratory IL-6; (E) pro-migratory IL-11; (F) pro-angiogenic VEGF; and (G) anti-angiogenic sEndoglin.

### 3.4. Inhibition of Let-7b-5p drives human first trimester trophoblast FLT1 expression in a NF**κ**B-dependent manner

To further explore the mechanism by which trophoblast sFlt-1 production is elevated under reduced Let-7b-5p conditions, similar to what is seen in preeclampsia [3, 6, 8, 9], we explored signaling pathways that are linked to the regulation of this anti-angiogenic factor production. When trophoblast Let-7b-5p expression was reduced, the phosphorylation status of p65 NFκB was significantly elevated by 4.3 ± 1.2-fold (Figure 4A & B). The activation of p38 MAPK, ERK and JNK pathways was not significantly altered when Let-7b-5p was inhibited (Figure 4A & B). When NFκB activity was inhibited under reduced Let-7b-5p conditions, trophoblast FLT1 mRNA expression was significantly reduced (Figure 4C).

**Figure 4.**
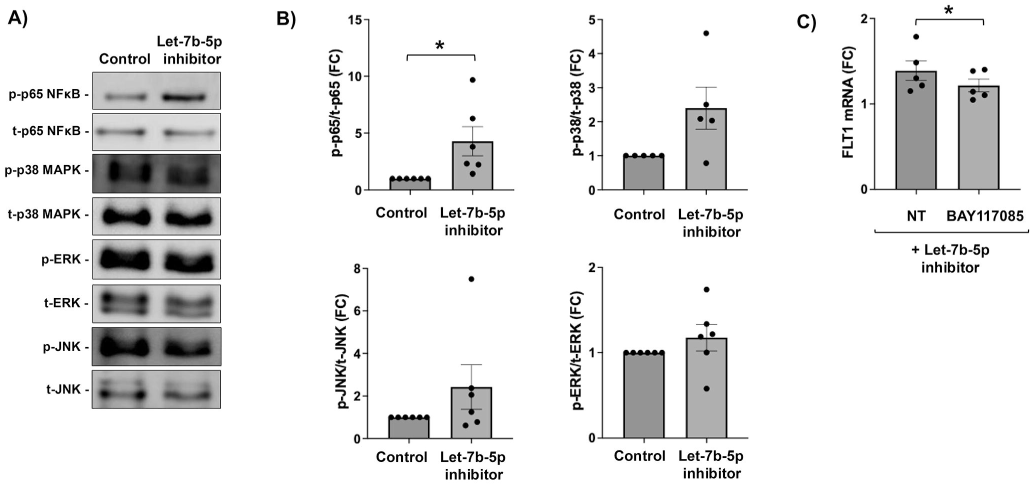
**Inhibition of Let-7b-5p drives human first trimester trophoblast FLT1 mRNA expression in a NF**κ**B-dependent manner.** (A-B) Trophoblast cells were transfected with a scramble control (control) or a Let-7b-5p inhibitor (n=6). After 72 hrs cell lysates were analyzed by Western blot for phosphorylated (p) and total (t) p65 NFκB, p38 MAPK, ERK and JNK. (A) Western blot images are from a representative experiment. (B) Bar charts show quantification of protein expression as determined by densitometry (\**p*<0.05; FC= fold change). (C) Trophoblast cells were transfected with a scramble control or a Let-7b-5p inhibitor and treated with no treatment (NT) or the NFκB inhibitor, BAY117085 (n=4). After 72 hrs, cellular RNA was measured for FLT1 mRNA levels by RT-qPCR using GAPDH as a housekeeping gene for normalization. Data is shown as the fold change (FC) of FLT1 relative to the scramble control (\**p*<0.05).

### 3.5. Inhibition of Let-7b-5p reduces trophoblast-endothelial interactions in a 3D Matrigel model

Reduced trophoblast migration, together with a pro-inflammatory and anti-angiogenic milieu results in reduced spiral arteriole remodeling, a seminal alteration that is considered pathognomonic of a preeclamptic placental phenotype [2–5]. We used an established 3D system that models trophoblast invasion into Matrigel towards pre-seeded endometrial endothelial cells and subsequent trophoblast-endothelial interactions, somewhat similar to that seen in spiral artery transformation [20]. In this system we found that reduced trophoblast Let-7b-5p expression significantly inhibited trophoblast invasion and subsequent trophoblast-HEEC interactions (Figure 5). In this model under control conditions, red HEECs form tube-like structures, resembling vessels in the Matrigel and when green trophoblast cells are introduced, they invade these tube-like structures, co-localize with, and replace, the endothelial cells. It is important to note that in the absence of trophoblast cells, the HEEC tubes formed in Matrigel will begin to destabilize after 48hrs [21]. After 4 hrs, when compared to the trophoblast cells transfected with a scramble control, trophoblasts transfected with the Let-7b-5p inhibitor showed less invasion into these endothelial tubes (Figure 5A) resulting in significantly less green fluorescence (Figure 5B). There were also significantly less red HEEC signals under trophoblast Let-7b-5p inhibitor conditions (Figure 5B), suggesting some active destabilization of the HEEC tubes when trophoblast Let-7b-5p expression was reduced.

**Figure 5.**
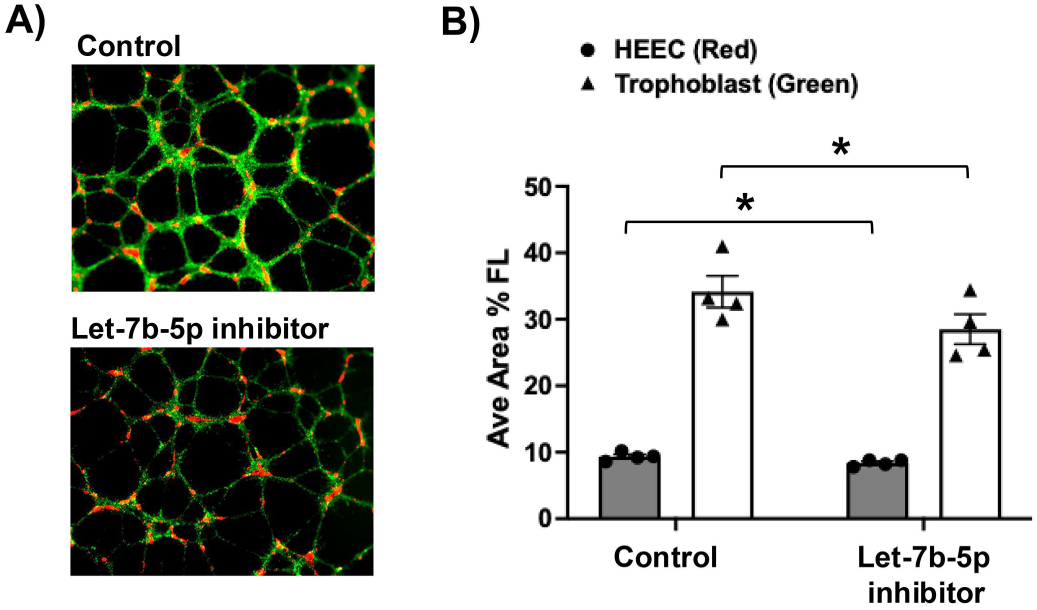
Inhibition of Let-7b-5p reduces trophoblast invasion and trophoblast-endothelial interactions in a 3D Matrigel model. (A-B) HEECs were stained red, seeded into Matrigel, and allowed to form tube. Trophoblast cells transfected with either a scramble control (control) or a Let-7b-5p inhibitor were stained green, seeded onto of the Matrigel and the co-culture system was incubated for 24 hrs (n=4). (A) Images of the co-culture are from a representative experiment. (B) Bar chart shows quantification of red and green fluorescence (n=4; \**p*<0.05).

### 3.6. Trophoblast sEVs with reduced Let-7b-5p inhibit HEEC angiogenesis

It is well established that trophoblast release small extracellular vesicles (sEVs) and use these to communicate with target cells, including uterine endothelial cells [14]. It has also been demonstrated that human first trimester trophoblast derived sEVs express Let-7b-5p [14]. We questioned whether reduced Let-7b-5p in trophoblast-derived sEVs might be one way in which rapid HEEC tube destabilization might be occurring. As shown in Figure 6A, transfection of trophoblast cells with a Let-7b-5p inhibitor significantly reduced trophoblast sEV expression of this miR by 89.9 ± 2.1%. Exposure of HEECs to control sEVs or Let-7b-5p inhibitor sEVs had no effect on HEEC cell viability (Figure 6B). However, using a 3D tube formation assay, as a measure of HEEC angiogenesis, Let-7b-5p inhibitor sEVs significantly reduced the ability of HEECs to form tubes by 24.3 ± 4.1%, when compared to control sEVs (Figure 6C).

**Figure 6.**
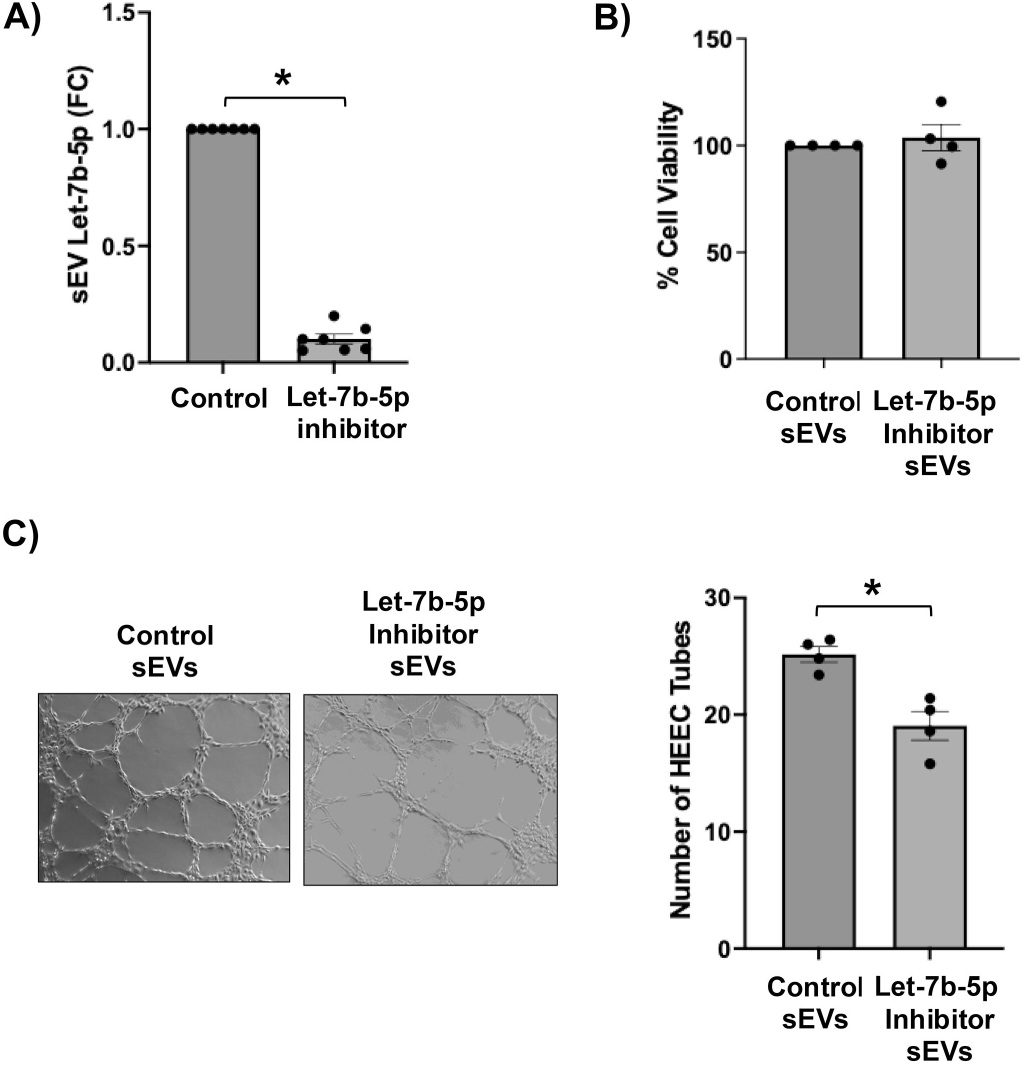
Trophoblast sEVs with reduced Let-7b-5p inhibit HEEC angiogenesis in a 3D tube formation assay. (A) sEVs were isolated from the culture supernatants of trophoblast cells transfected with either a scramble control or a Let-7b-5p inhibitor. Following isolation, trophoblast sEV RNA was measured for Let-7b-5p by RT-qPCR and expressed as fold change (FC) relative to the control (n=7; \**p*<0.05). (B-C) HEECs were exposed to sEVs isolated from trophoblast cells transfected with either a scramble control (Control sEVs) or a Let-7b-5p inhibitor (Let-7b-5p Inhibitor sEVs) at 1x10^9^/ml. (B) After 24 hrs, HEEC cell viability was measured (n=3). (C) After 4 hrs HEEC tube formation in Matrigel was imaged. Images are from a representative experiment and the bar chart show quantification of HEEV tubes (n=4; \**p*<0.05).

## 4. Discussion

Preeclampsia is a common hypertensive disorder of pregnancy characterized by a pro-inflammatory, anti-migratory and anti-angiogenic placental phenotype. The pathological underpinnings are thought to be established early in gestation, stemming from trophoblast dysfunction [2–5]. However, little is known about what mechanisms govern normal trophoblast function or what changes occur that predispose pregnant individuals to the risk of developing preeclampsia later in the course of pregnancy. Herein, we report that the miR, Let-7b-5p, drives normal human first trimester trophoblast migration through activation of TLR7 and TLR8, and in a TLR7-dependent manner, this miR negatively regulates trophoblast anti-angiogenic sFlt-1 production. Thus, Let-7b-5p may play a functional role in normal placentation. We further demonstrate that inhibition of trophoblast Let-7b-5p reduced trophoblast migration, elevated trophoblast release of sFlt-1, and reduced trophoblast invasion and subsequent endometrial endothelial interactions. Thus, reduced trophoblast Let-7b-5p appears to promote a pathological phenotype similar to that seen in preeclampsia.

First trimester trophoblast migration and invasion are reduced during preeclampsia [24, 25], however, little is known about the mechanisms behind this functional change. Human placental Let-7b-5p levels are higher in the third trimester when compared to the first trimester [7], but there is a reduction in Let-7b-5p expression in the placentae and circulation of women with preeclampsia [6, 8, 9], suggesting that alterations in this miR may have a pathophysiological impact. Indeed, in human first trimester trophoblast cells, Let-7b-5p overexpression was previously shown to positively drive cell invasion by activating the ERK signaling pathway through suppression of TGFBR1 [6]. Since Let-7b-5p is a member of a small family of miRs that can function non-classically by activating TLR7 and TLR8, we questioned whether this signaling pathway might also be involved in its regulation of trophoblast function. Similar to the study by Gao *et al.*, [6], we found that overexpression of Let-7b-5p elevated human first trimester trophoblast cell migration without affecting cell viability/proliferation. Furthermore, we found that Let-7b-5p was driving spontaneous trophoblast migration through activation of TLR7 and TLR8, highlighting its function as a TLR7/TLR8-activating miR. We previously found a role for another TLR7/TLR8-activating miR, miR-146a-3p, in driving human first trimester trophoblast inflammatory IL-1β and IL-8 production in a model of obstetric antiphospholipid syndrome [11, 12], and we found a role for miR-146a-3p in mediating human fetal membrane inflammation in models of infection [15, 16, 26]. However, our study indicates that this Let-7b-5p-TLR7/TLR8 signaling pathway may be important in normal trophoblast physiology. Additionally and conversely, inhibition of basally expressed Let-7b-5p reduced trophoblast cell migration, without affecting cell viability, aligning with the observed reduced Let-7b-5p levels in women with preeclampsia [6, 8, 9].

Using the HTR8 trophoblast cell line we previously demonstrated that endogenous IL-6 contributes to trophoblast migration through activation of the STAT3 pathway [27]. Gao *et al.*, reported that Let-7b-5p overexpression in these HTR8 cells elevated IL-6 and reduced TNFα at the mRNA level, but they did not show this at the protein level, nor did they test STAT3 activity [6]. In our studies, using the Sw.71 trophoblast cell line, while Let-7b-5p modulation regulated trophoblast migration, it did not influence the pro-migratory factors IL-6 or IL-11 [27, 28] at the protein level, indicating that downstream of TLR7/TLR8, activation an alternative mechanism of migratory regulation is involved. Additionally, trophoblast IL-1β, IL-8 and TNFα protein secretion were unaffected when Let-7b-5p expression was modulated, indicating that Let-7b-5p activation of TLR7 and TLR8 results in distinct downstream signaling and cellular function when compared to miR-146a-3p activation of these TLRs in the trophoblast [11, 12].

In addition to reduced trophoblast migration and invasion, in preeclampsia, there is also an increase in anti-angiogenic factors [29–31]. Indeed, elevated anti-angiogenic sFlt-1 late in gestation can be predictive for a woman’s risk for preeclampsia [31]. During normal placentation, pro-angiogenic factors, such as VEGF and PlGF promote the remodeling of spiral arterioles, resulting in an increase in blood flow. However, in preeclamptic pregnancies, elevated sFlt-1 inhibits the function of VEGF and PlGF, resulting in reduced vascular remodeling, and a hypertensive environment [4, 31–33]. The mechanism underlying elevated sFlt-1 is not known. In this current study, we found that inhibition of basally expressed Let-7b-5p elevated trophoblast sFlt-1 release at the mRNA and protein level. In contrast, Let-7b-5p overexpression blunted trophoblast sFlt-1 production. This suggests that under normal conditions, endogenous Let-7b-5p serves to negatively regulate trophoblast sFlt-1, and when Let-7b-5p expression is reduced, as it is in preeclampsia, this brake is released, allowing elevated sFlt-1 release. The mechanism by which Let-7b-5p negatively regulates trophoblast sFlt-1 was found to be through activation of TLR7, but not TLR8, indicating that these receptors both can be triggered by Let-7b-5p but can mediate distinct responses. Furthermore, our findings indicate that when Let-7b-5p is inhibited this brake is removed, allowing expression of an alternate splice variant of trophoblast FLT1 (V2) via NFκB that gives rise to elevated sFlt-1 protein [34]. Thus, we found that Let-7b-5p regulates trophoblast sFlt-1 at the mRNA level, rather than regulating its release at the protein level by MMP14-mediated proteolytic cleavage [23], which we found to be unchanged. While pro-angiogenic VEGF expression is reduced in preeclamptic pregnancies, and the anti-angiogenic factor sEndoglin is elevated [30, 32], we found that in the first trimester trophoblast cells, Let-7b-5p did not influence trophoblast secretion of these angiogenic factors suggesting that unlike sFlt-1, the mechanism regulating VEGF and sEndoglin is independent of Let-7b-5p. Similarly, we were not able to detect PlGF secretion by the Sw.71 cells line.

Appropriate trophoblast migration and the angiogenic balance is important for promoting normal trophoblast invasion and remodeling of the uterine spiral arterioles, and thus for normal placentation. Disruption in the crosstalk between the trophoblast and the uterine vascular endothelium is common in preeclampsia and this manifests as shallow trophoblast invasion and reduced spiral artery transformation. Using an *in vitro* 3D system that models trophoblast invasion towards endometrial endothelial cells and subsequent trophoblast-endothelial interactions, somewhat similar to that seen in spiral artery transformation [20], we found that inhibition of trophoblast Let7b-5p expression reduces the normal interactions between the trophoblast and endothelial tube-like structures. Moreover, this disrupted trophoblast-endothelial crosstalk and associated endothelial angiogenesis may be mediated by trophoblast-derived sEVs lacking Let-7b-5p. Together these observations underscore the importance of Let-7b-5p in regulating normal trophoblast function and placentation.

One limitation of this study is the use of cell lines rather than primary cells. Obtaining any human first trimester trophoblast cells is extremely difficult. Moreover, performing transfections and highly mechanistic studies, as presented herein, requires large cell numbers that can be cultured robustly for long periods of time. Thus, the use of primary cells for such experiments poses extreme challenges. Lastly, while we utilized 3D culture systems to model angiogenesis and trophoblast-endothelial interactions, we acknowledge that these experimental approaches are limited, that the processes of placentation and spiral artery transformation is more complex, and that our finds are not direct evidence of the *in vivo* scenario.

In summary, we report that human first trimester trophoblast migration is positively driven by Let-7b-5p activating TLR7 and TLR8, while in a TLR7-dependent manner, Let-7b-5p negatively regulates trophoblast anti-angiogenic sFlt-1 production. Inhibition of trophoblast Let-7b-5p reduces trophoblast migration and normal interactions with endometrial endothelial cells, while sFlt-1 production is elevated. Together, this work highlights a role for TLR7/TLR8-activating Let-7b-5p in promoting normal trophoblast function and endothelial interactions and that disruption in this miR-driven signaling pathway may be, in part, causative to the development of a preeclamptic placental phenotype.

## CRediT Authorship Contribution Statement

Emily Siegel: Investigation, Writing – original draft

Lily Salmeron: Investigation, Writing – review & editing

Vikki Abrahams: Conceptualization, Supervision, Formal Analysis, Writing – review & editing

Lubna Pal: Conceptualization, Supervision, Funding acquisition, Project administration, Writing – review & editing

## Declaration of Completing Interest

The authors have nothing to disclose.

